# Synthesizing developmental trajectories

**DOI:** 10.1101/157834

**Authors:** Paul Villoutreix, Joakim Andén, Bomyi Lim, Hang Lu, Ioannis G. Kevrekidis, Amit Singer, Stanislav Y. Shvartsman

## Abstract

Dynamical processes in biology are studied using an ever-increasing number of techniques, each of which brings out unique features of the system. One of the current challenges is to develop systematic approaches for fusing heterogeneous datasets into an integrated view of multivariable dynamics. We demonstrate that heterogeneous data fusion can be successfully implemented within a semi-supervised learning framework that exploits the intrinsic geometry of high-dimensional datasets. We illustrate our approach using a dataset from studies of pattern formation in Drosophila. The result is a continuous trajectory that reveals the joint dynamics of gene expression, subcellular protein localization, protein phosphorylation, and tissue morphogenesis. Our approach can be readily adapted to other imaging modalities and forms a starting point for further steps of data analytics and modeling of biological dynamics.

## 1. Introduction

The need to synthesize data from different observations into coherent multivariable trajectories is discussed in multiple contexts, from physics to social sciences, but systematic approaches for accomplishing this task have yet to be established (1–5). Here we address this task for imaging studies of developing tissues, where patterns of cell fates are established by complex regulatory networks (6–8). Advances in live imaging continue to provide new insights into the dynamics of individual components in these networks, but imaging more than three reporters at the same time is still challenging and limited to model genetic organisms (9, 10). Furthermore, in the absence of reliable live reporters, dynamics of some state variables can only be inferred from fixed tissues. Because of these limitations, extracting the multivariable dynamics from the heterogeneous datasets collected by imaging of live and fixed tissues becomes a non-trivial task (11, 12).

The problem can be illustrated by an imaging dataset from the early Drosophila embryo (Fig. 1A-B), a model system in which a graded profile of the nuclear localization of transcription factor Dorsal (Dl) establishes the dorsoventral (DV) stripes of gene expression that control cell fates and tissue deformations (13–15). Current mechanisms of the DV patterning system invoke multiple state variables, such as the levels of gene expression and protein phosphorylation (16) (Fig. 1C). These mechanisms were elucidated in studies that reveal only a small subset of the full state space, most commonly 2-3 variables per experiment. Can these partial views be fused into a consistent multivariable trajectory? This is a general question that applies to essentially all developmental systems.

**Fig. 1.**
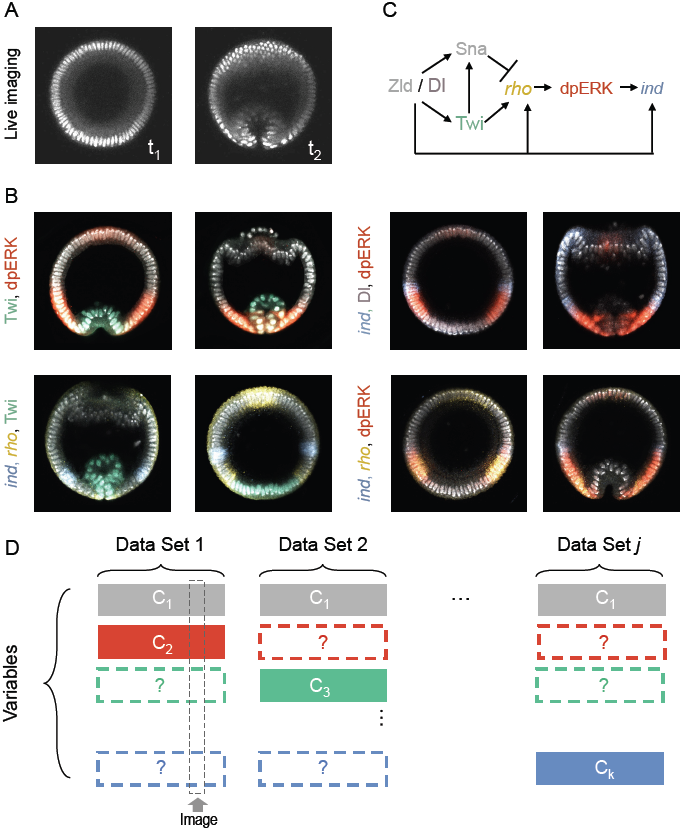
Stating the problem of data fusion. (A-B) Example datasets of molecular signals and morphology during the DV patterning of Drosophila embryo; all images are collected from optical cross-section along the DV axis, ~ 15% from the posterior pole of the embryo. (A) Frames from a live imaging movie, showing positions of nuclei during the early stages of gastrulation. (B) Images of fixed embryos, stained with probes and antibodies revealing the spatial patterns of nuclear Dl (pink), Twi (green), dually phosphorylated ERK (red), and transcripts of rho (yellow) and ind (blue). (C) A fragment of a DV patterning network in the early Drosophila embryo. (D) Data fusion as a matrix completion problem: Each row corresponds to a variable, e.g. nuclear positions, gene expression levels, time stamp, revealed by visualizing different molecular or cellular components, nuclei, transcripts, or protein phosphorylation. Each column of the matrix corresponds to an image giving access to some of the states through various channels. The remaining states, labeled with a question mark, must be estimated from other datasets.

We realized that this question can be addressed by casting the task of data fusion as a matrix completion problem (Fig. 1D). Specifically, an image of a fixed embryo or a frame from a live imaging movie can be viewed as a column in a matrix where rows correspond to the relevant variables, such as developmental time or the level of gene expression at a given position. Because of limitations in the number of states that can be accessed simultaneously, the matrix is incomplete. For example, live imaging of gastrulation provides information about nuclear positions as a function of time, but is silent about the levels of gene expression. On the other hand, an image of a fixed embryo reveals the distribution of an active enzyme but has no direct temporal information. Thus, multivariable data fusion requires completing this matrix, filling in the missing components by estimates informed by the rest of the data. Below we show how this task can be accomplished by solving a suitably posed semi-supervised learning problem. We first provide a closed-form solution to this problem and then demonstrate its successful performance on synthetic and experimental datasets.

## 2. Results

### Semi-supervised learning framework for matrix completion

We assume here that all experiments contain a common variable, which is sufficient to determine all other variables that can be measured or to be predicted. For instance, this variable is revealed by a signal that reports positions of nuclei. This means that the first row in the matrix are complete. To complete other rows, we must establish the mappings between the common variable and each of the target variables. These mappings can be found within a semi-supervised learning framework, in which the values of the variables in the incomplete rows are estimated from a training dataset (17, 18).

As an example, consider images from fixed embryos that are stained to reveal the spatial pattern of an active enzyme, visualized using a phosphospecific antibody (Fig. 2). They provide labeled data points that contain information about the common variable and a specific target variable. On the other hand, images without this staining, such as the frames from live imaging of morphogenesis, provide unlabeled data points with only the common variable. By finding a mapping between the common and target variables, we can essentially “color” the frames of a live imaging movie by snapshots of molecular patterns from fixed embryos.

**Fig. 2.**
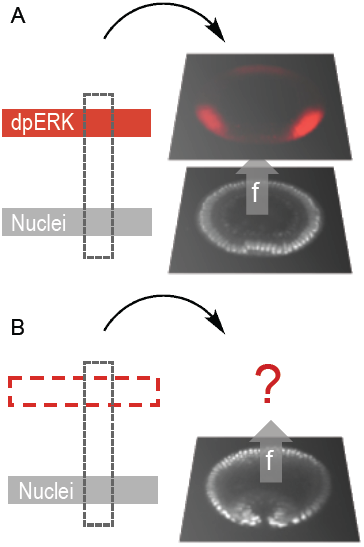
Learning a mapping from a common channel. An experimental image can be decomposed into various channels. E.g., red: dpERK, visualized with a phosphospecific antibody, gray: nuclei, visualized through either DAPI (in fixed images) or Histone-RFP (in live imaging). The training ensemble of labeled images (A) is used to predict the labels on a set of unlabeled images (B) using common information, the morphology obtained through the nuclei signal in this case. Morphological proximity yields similar labels.

A critical assumption in finding the mappings is that the multivariable dynamics of the patterning process are both low-dimensional and smooth. This assumption is supported by studies with mathematical models of specific biological systems and by computational analysis of datasets from imaging studies of development (19, 20). More formally, we consider a set of data points (*x*_1_*, …, x*_*l*_*, x*_*l*+1_*, …, x*_*l*+*u*_) belonging to a space χ. These points correspond to the values of the common variable in the complete row. On the other hand, a row corresponding to any one of the target variables is incomplete. The values in the filled columns of this row are called labels. These are denoted (*y*_1_*, …, y*_*l*_) and belong to a target space *Y*. The semi-supervised learning techniques transfer the information contained in the labeled data points ((*x*_1_*, y*_1_)*, …,* (*x*_*l*_*, y*_*l*_)) to the unlabeled points (*x*_*l*+1_*, …, x*_*l*+*u*_), while preserving the intrinsic structure of the dataset(18). Stated otherwise, these techniques *learn* the mapping *y* = *f* (*x*) assuming that the considered process is smooth, which means that similar values of *x* give rise to similar values of *f* (*x*).

The missing values, corresponding to the unlabeled data points in each of the incomplete rows, are found by solving the following optimization problem:

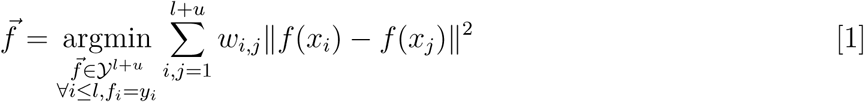

where 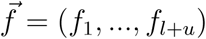 are the values of the target variable on the data points (*x*_1_*, …, x*_*l*+*u*_), the considered norm in *Y* is the Euclidean distance and the weights *w*_*i,j*_ represent the similarity between two data points *x*_*i*_ and *x*_*j*_. The norm in the space χ can for example be the Euclidean distance in the space where each dimension corresponds to an image pixel or some coordinate in an arbitrary feature transform of that image.

This quadratic optimization problem, known as harmonic extension, has a unique solution that relates the unlabeled data points *f*_*l*+1_*, …, f*_*l*+*u*_ to the labels *y*_1_*, …, y*_*l*_ where *Y* = ℝ (17, 21). The explicit solution reads:

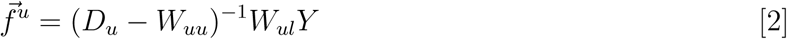

where *Y* = (*y*_1_*, …, y*_*l*_) and 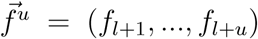, and 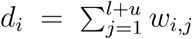, and *D*_*u*_ = diag(*d*_*l*+1_*, …, d*_*l*+*u*_), *W*_*uu*_ = (*w*_*i,j*_)_*l*+1*≤i,j≤l*+*u*_, and 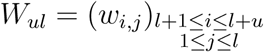 (*Methods and Materials*).

### Illustrative example

To illustrate our method, we considered a one-dimensional nonlinear trajectory in a three-dimensional space. The trajectory is given by the set of equations

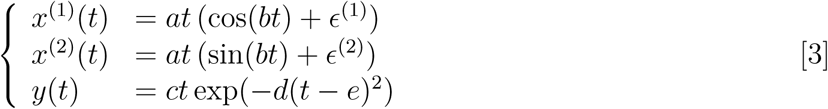

where *a, b, c, d, e* are constants, *ϵ* ^(1)^ and *ϵ*^2)^ are Gaussian noise sources and *t* is a real-valued parameter. The set of points (*x*^(1)^(*t*)*, x*^(2)^(*t*)) forms a one-dimensional non-linear manifold embedded in the two dimensional plane and it is parameterized by *t*. These points are analogs of the embryo morphology. In the absence of noise, this mapping from *t* to the 2D plane can be inverted as 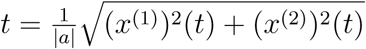. The signal *y*(*t*) is a smooth function of *t* and is thus a smooth function of (*x*^(1)^*, x*^(2)^) by composition. In this example, *y* corresponds to the target modality that we would like to estimate.

To mimic the setting of data fusion with three modalities, ((*x*^(1)^*, x*^(2)^)*, t, y*), we consider the following situation: suppose that one acquire a set of labeled points, i.e. a set of *l* triplets, (((*x*^(1)^(*t*_1_)*, x*^(2)^(*t*_1_))*, y*(*t*_1_))*, …,* ((*x*^(1)^(*t*_*l*_)*, x*^(2)^(*t*_*l*_))*, y*(*t*_*l*_))) and a set of *u* unlabeled, but timestamped, points, (((*x*^(1)^(*t*_*l*+1_)*, x*^(2)^(*t*_*l*+1_))*, t*_*l*+1_)*, …,* ((*x*^(1)^(*t*_*l*+*u*_)*, x*^(2)^(*t*_*l*+*u*_))*, t*_*l*+*u*_)), as shown in Fig. 3A-B. The pairwise similarity measures *w*_*i,j*_ are computed using Euclidean norm between pairs of data points (*x*^(1)^(*t*_*i*_)*, x*^(2)^(*t*_*i*_)) and (*x*^(1)^(*t*_*j*_)*, x*^(2)^(*t*_*j*_)). Then, using Equation (2) it is possible to estimate *y* = *f* (*x*) on the set of unlabeled data points using the harmonic extension algorithm. The results are shown in Fig. 3C. We then directly obtain *y* as a function of *t* by composition using the known time stamps (*t*_*l*+1_*, …, t*_*l*+*u*_).

**Fig. 3.**
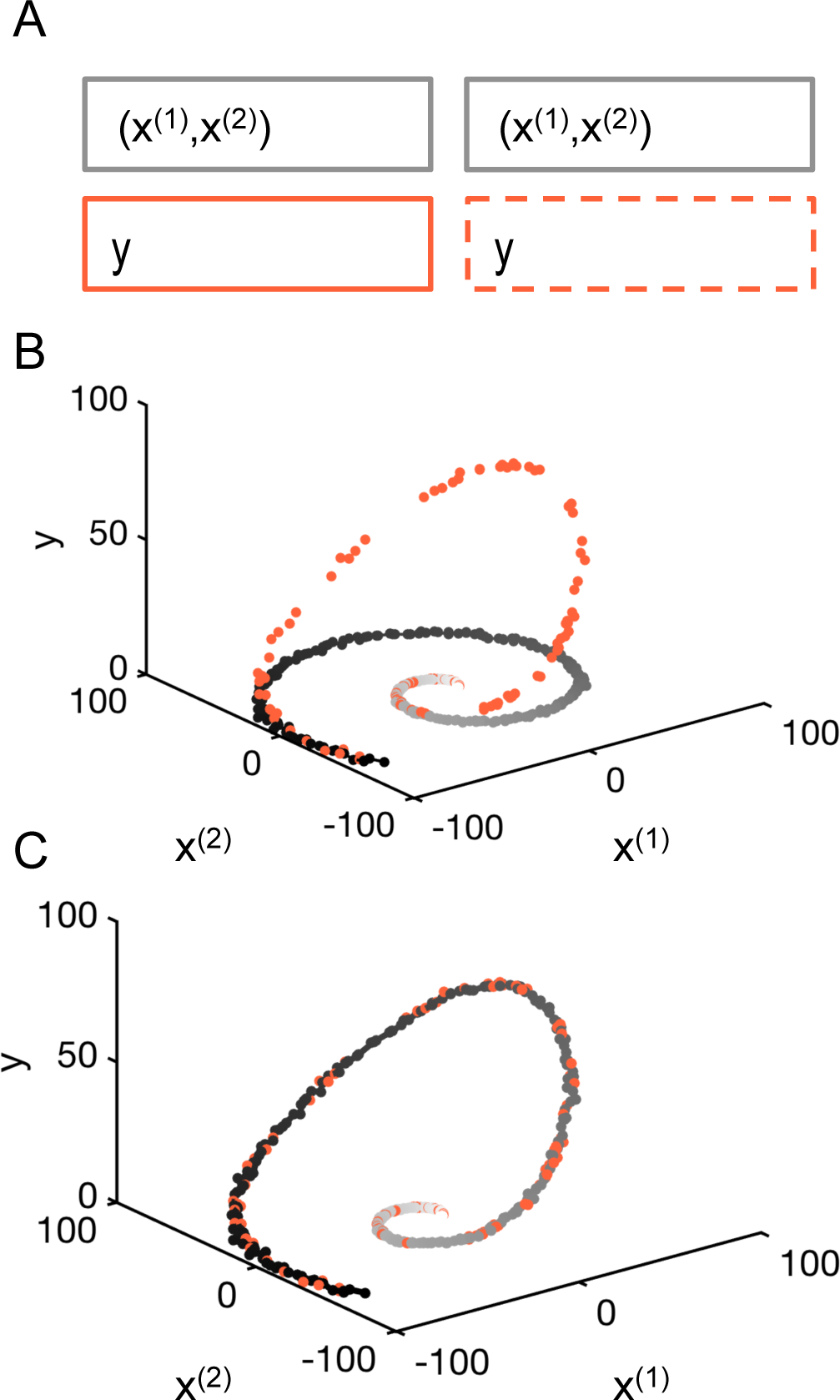
Illustrative example. (A) Matrix formulation of the problem with 120 labeled samples, (((*x*^(1)^(*t*_1_)*, x*^(2)^(*t*_1_))*, y*(*t*_1_))*, …,* ((*x*^(1)^(*t*_*l*_)*, x*^(2)^(*t*_*l*_))*, y*(*t*_*l*_))), and 300 unlabeled samples (((*x*^(1)^(*t*_*l*+1_)*, x*^(2)^(*t*_*l*+1_))*, t*_*l*+1_)*, …,* ((*x*^(1)^(*t*_*l*+*u*_)*, x*^(2)^(*t*_*l*+*u*_))*, t*_*l*+*u*_)). (B) The points are distributed on a non-linear 1-dimensional manifold in the -plane. Some points, the snapshots, contain a value for the signal. (C) Result of the interpolation on the nonlinear manifold using the harmonic extension algorithm.

The accuracy of the estimated multivariable dynamics can be assessed using a K-fold validation strategy on the labeled samples (Fig. S1 and *Methods and Materials*). For chosen set of parameters and the size of the dataset, the error is *∼*1%. As expected for the semi-supervised learning framework, the error decreases with the addition of new unlabeled data points. This example demonstrates how the proposed approach successfully recovers multivariable dynamics from heterogeneous datasets that combine continuous views for part of the state variables and snapshots that report several states without direct temporal information.

### Fusion of imaging datasets

As a representative dataset from imaging studies of multivariable dynamics in living systems, we use a collection of *∼*1000 images each of which reveals the spatial position of the nuclei and either a timestamp or the distribution of one or several components of the DV patterning network (Fig. 1D). To apply the semi-supervised learning approach to data fusion to this dataset we need to compute pairwise similarities between the images using the common channel. Prior to this, we took several preprocessing steps that aim to minimize image variability associated to sample handling, microscope calibration and imaging. First, the images were registered to align their ventral-most points. The images were then resized and cropped such that the embryos occupy 80% of the image. All images were resized to 100 by 100 pixels. To overcome local variations of image intensity, we computed a local average using a Gaussian kernel, and then renormalized the image by that value. We also applied a logistic function to the images to handle contrast variability, Fig. S2.

Most importantly, to ensure that pairwise differences between images are insensitive to small translations or deformations, we applied the scattering transform (22) and compared the resulting transform vectors. The scattering transform of an image is a signal representation obtained by alternating wavelet decompositions and pointwise modulus operators. We found that second-order scattering coefficients with an averaging scale of 64 pixels provided sufficient invariance. These are computed using the ScatNet toolbox (23, 24). The result is a vector of dimension 784 for each image. The point clouds corresponding to each of the 11 datasets were centered separately. The corresponding low-dimensional manifold on which the data points lie is shown on Fig. S4.

For each of the 512x512 pixels of each live movie frames, there is a common channel reporting the nuclei spatial position and there are 5 channels that we would like to complete. These channels contain the information about the spatial distributions of one enzyme (dpERK), two transcription factors (Twist and Dorsal), and transcripts of two genes (ind and rho). We thus solved the data fusion problem for each pixel and each channel, leading to 5x512x512 semi-supervised learning solutions. The combination of labeled and unlabeled datasets is described on Table S2. The result is a multivariable trajectory for the joint dynamics of tissue shape and five molecular components within the regulatory network that patterns the DV axis of the embryo (Fig. 4). To evaluate the accuracy of the method, we computed the cross-validation error for each pixel and averaged over the entire images. We found that the normalized absolute error is of 0.9-2.5% of the signal range when considering the various modalities of the entire experimental datasets (Table S3). We show how the algorithm performs on several examples in Fig. 5.

**Fig. 4.**
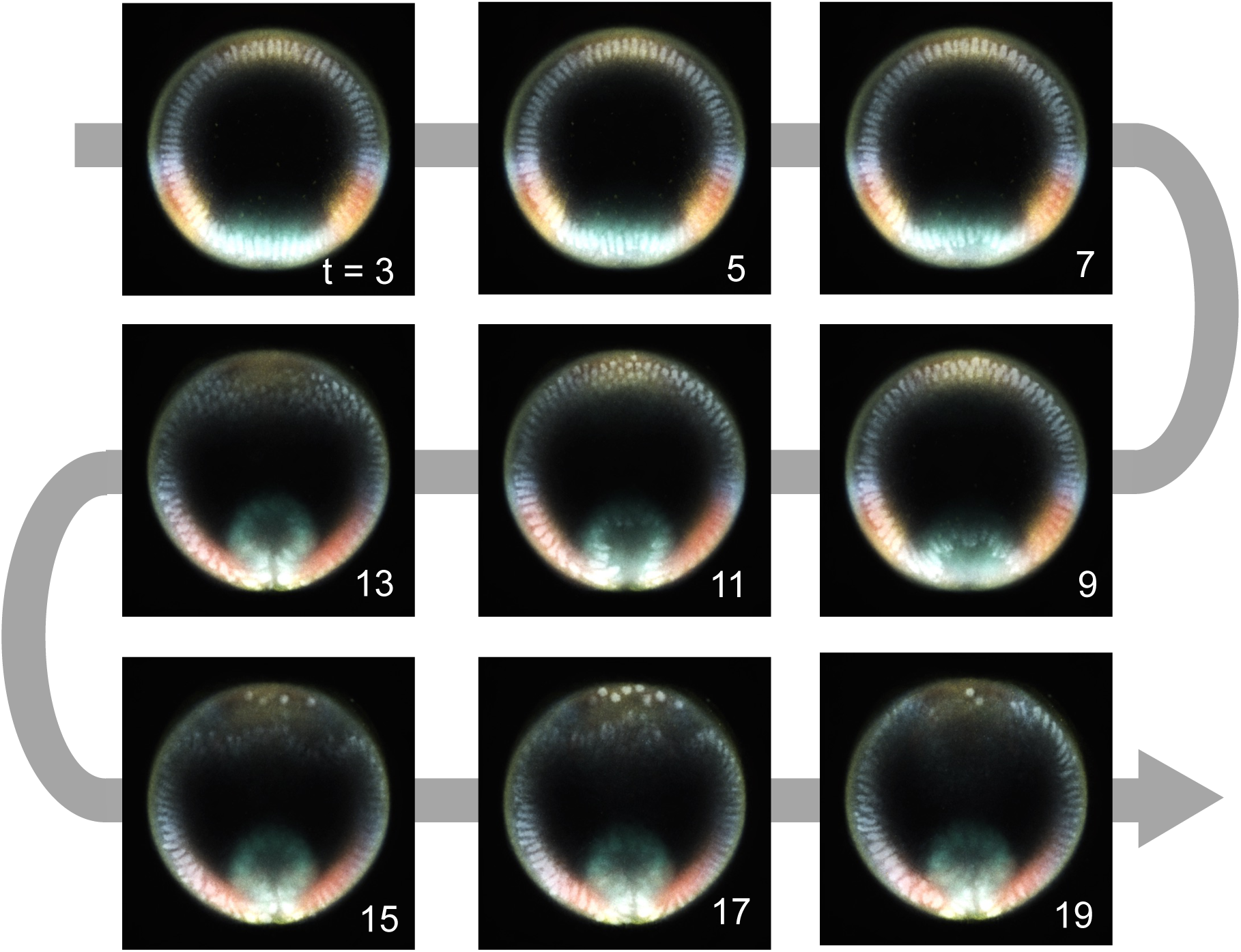
Colored movie frames obtained with our data fusion algorithm. The temporal resolution is 2 min, extracted from a 30 s resolution movie. The time stamps on the images are in min and indicate elapsed time from the start of the live movie. The colors correspond to dpERK (red), Dl (pink), rho (yellow), ind (blue), Twi (green).

**Fig. 5.**
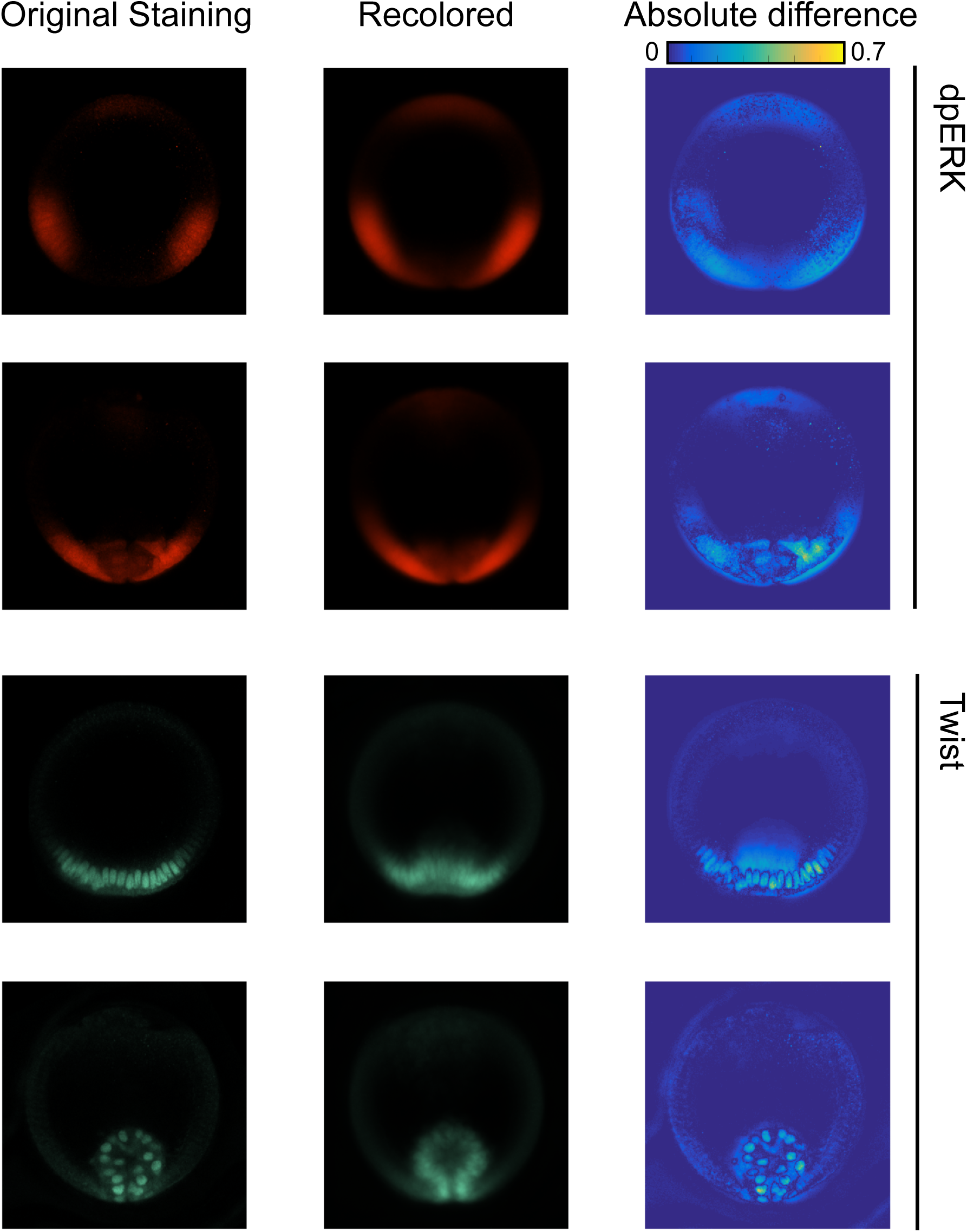
Examples showing how the algorithm performs on four fixed samples. The first column shows the original measurement, the second column shows the result of recoloring the snapshot through K-fold cross validation, and the third column shows the absolute difference between the original and recolored images, normalized by the signal range.

### Discussion

To summarize, we presented a formal approach to synthesizing developmental trajectories. By posing the task of data fusion as a semi-supervised learning problem, we obtained a closed form expression for the estimated values of all variables using harmonic extension. The reconstructed trajectories provide the basis for the more advanced mechanistic studies of multivariable processes responsible for the highly reproducible dynamics of developmental pattern formation. Our approach can also be extended to other semi-supervised learning methods (25), if the dimensionality of the intrinsic geometry is greater than one or if there is no unique common channel among all experiments. Most of the previous attempts to accomplishing this task explored specific features of developmental systems, such as the expression level of a particular gene, and used a discrete number of temporal classes, usually defined in ad hoc way (16, 26). Our approach reconstructs continuous time dynamics and relies on the intrinsic geometry of multidimensional datasets.

We conclude by pointing two directions for the future extensions and applications of the presented approach. First, while there are no conceptual limitations in using the presented matrix completion framework to studies of pattern formation and morphogenesis problems in three-dimensions(27), it is important to increase the computational efficiency of our approach, which can be done at multiple levels, starting with dimensionality reduction at the preprocessing step. At the same time, for a large class of patterning processes that happen on the surfaces of epithelial sheets, one can use the recently developed “tissue cartography” approach to first flatten the three-dimensional images (5), which should make our approach directly applicable. Second, following the step of data fusion, one can attempt to model the observed multivariable dynamics. Here one can employ several modeling methodologies, from mechanistic modeling of specific molecular and tissue-level processes (28–32), to equation-free approaches, which aim to deduce the underlying mechanisms directly from data (33, 34). Going beyond working with already existing datasets, it will be important to establish an adaptive data assimilation framework, whereby data collection itself is guided by the future need to reconstruct the multivariable trajectories. These ideas are already being discussed in the context of transcriptional profiling studies of cell differentiation and can be extended to synthesis of multivariable trajectories from imaging studies (35).

## Methods and Materials

Extended methods and materials are presented in *SI Appendix*.

### Image datasets

All images are cross-sections of Drosophila embryos taken at *∼*90*μ*m from the posterior pole. Time-lapse movies were obtained using a Nikon A1-RS confocal microscope with a 60x Plan-Apo oil objective. The nuclei were stained with Histone-RFP. A total of 7 movies was acquired with a time resolution of 30 seconds per frame. All movies start about 2.5 hr after fertilization and end after about 20 min after gastrulation starts (about 3.3 hr after fertilization). Four datasets of fixed images were acquired to visualize nuclei, protein expression of dpERK, Twist, and Dorsal, and mRNA expression of ind and rho. Immunostaining and fluorescent in situ hybridization protocols were used as described before13. DAPI (1:10,000; Vector laboratories) was used to visualize nuclei. Rabbit anti-dpERK (1:100; Cell Signaling), mouse anti-Dorsal (1:100; DSHB), rat anti-Twist (1:1000; gift from Eric Wieschaus, Princeton University), sheep anti-digoxigenin (1:125; Roche), and mouse anti-biotin (1:125; Jackson Immunoresearch) were used as primary antibodies. Alexa Fluor conjugates (1:500; Invitrogen) were used as secondary antibodies. Stained embryos were imaged using Nikon A1-RS confocal microscope with a 60x Plan-Apo oil objective. Embryos were mounted in a microfluidic device for end-on imaging, as described previously (16, 36). The first dataset contains 108 images stained with rabbit anti-dpERK and rat anti-Twist antibodies. The second dataset contains 59 images stained with mouse anti-Dorsal antibody, rabbit anti-dpERK antibody, and ind-DIG probe. The third dataset contains 58 images stained with ind-biotin probe, rho-DIG probe, and rabbit-dpERK antibody. The fourth dataset contains 30 images stained with rat anti-Twist antibody, ind-biotin probe, and rho-DIG probe. The distribution of the datasets as labeled and unlabeled data depending on the considered variable is summarized on Table S2. Raw images are can be found in Supplementary Files 1 or on the public github repository https://github.com/paulvill/data-fusion-images.

### The affinity matrix

The affinity matrix *W* = (*w*_*i,j*_) is computed using a Gaussian kernel *w*_*i,j*_ = 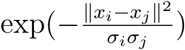 with scaling parameters *σ* and *σ* computed locally as the average of the distance with respect to the 10 closest neighbors as described in SI appendix. We used the Euclidean norm in the space of scattering transformed data. The resulting affinity matrix is shown on Fig. S3. The corresponding underlying one-dimensional manifold is shown on Fig. S4.

### Computing the cross validation error

The K-fold cross validation error was computed by extracting subsamples of the labeled data points and the semi-supervised learning framework was used to predict the value of the labels on them. For the image datasets, we computed the absolute error between the actual value of pixel intensity to the predicted one. The absolute error was then normalized by the range of the signal computed from the entire set of images for a given channel. The number of bins K was chosen so that the number artificially unlabeled data points was about 20. The results for each dataset are shown in Table S3 and described in SI appendix.

### Movie coloring

The result of data fusion led to multimodal time lapses of developing embryo showing nuclei and the spatio-temporal dynamics of dpERK, Dl, rho, ind, and Twi. The images were colored using the color code shown in Table S4, i.e. dpERK (red), Dl (pink), rho (yellow), ind (blue), Twi (green). A resulting colored movie is provided in Supplementary Files 2.

### Code implementation

The semi-supervised framework used to accomplish the task of data fusion is completely implemented in the open-source MATLAB library and fully runs in GNU Octave. It is available as Supplementary Software or on the public github repository https://github.com/paulvill/data-fusion.

## Acknowledgments

PV was partially supported by the WIN program between Princeton University and the Weizmann Institute. SYS and YGK were supported by the NSF award MCB 1516970. YGK and AS were supported by NSF award CMMI-1310173. JA and AS are supported by Simons Investigator Award and Simons Collaboration on Algorithms and Geometry from Simons Foundation, and the Moore Foundation Data-Driven Discovery Investigator Award to AS. HL was supported by the GM088333 award from the NIH. We thank Carmeline Dsilva, Mahim Misra, Roy Lederman and David Sroczynski for helpful discussions.

## 1. Semi-supervised learning

### Problem Setting

We define the problem of data fusion using semi-supervised learning in general terms. A classical setting is to have a modality *x* belonging to some space χ which is common to all experiments and *M* other modalities (*y*^(1)^*, …, y*^(*M*)^) that vary across experiments. The difficulty here is that only a certain number of modalities can be measured simultaneously, so our goal is to fill in the missing modalities. The critical assumption that we use is that the common modality *x* is sufficient to determine the other modalities. In other words, there exists a set of functions 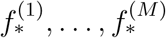 such that 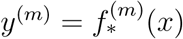 for all 1 ≤ *m* ≤ *M* and all *x*.

Let us represent the *i*th experiment, here containing the common modality *x* and the modalities *m*_1_, and *m*_2_ as an example by

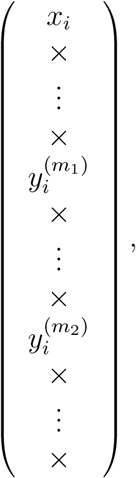

where × denotes an empty, or unmeasured, modality. The corresponding completed vector would then be

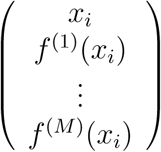

where *f* ^(1)^*, …, f* ^(*M*)^ are our estimates of the underlying functions 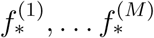 with 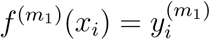 and 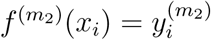.

### Problem formulation

The problem of fusing multiple modalities can be summarized as completing the following matrix using the information on the row *x* while preserving the continuity of the underlying manifold.

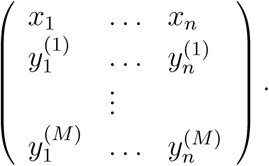

For the *m*th modality, we denote the set of experiments where it is measured by Ω(*m*). This is a subset of *{*1*, …, n}*, in which it has the complement 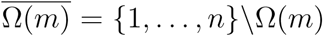. We shall refer to Ω(*m*) as the set of labeled datapoints, while 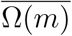 is the set of unlabeled data points for the *m*th modality. Note that we do not impose any particular structure on the sets of labeled points Ω(*m*). They can occur in any order and vary in size between the different modalities.

### Multiple semi-supervised learning by harmonic extension

The problem of completing the *m*th row of the matrix using the measured elements 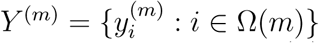 can be posed as a semi-supervised learning problem, which consists in learning the mapping *f* ^(*m*)^(*x*) = *y*^(*m*)^ on each of the points (*x*_1_*, …, x*_*n*_) (1). To simplify the notation, we shall consider the *n* values of *f* ^(*m*)^ on these points as a column vector in ℝ^*n*^. In addition, we denote the subset of its values when restricted to Ω(*m*) and 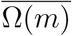 by *f* ^(*m*)^*|*_Ω(*m*)_ *∈* ℝ^*|*Ω(*m*)*|*^ and 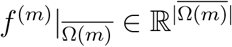, respectively, where *|A|* denotes the number of elements in the set *A*. Since *Y* ^(*m*)^ is also a column vector in ℝ^*|*Ω(*m*)*|*^, we formulate our problem as

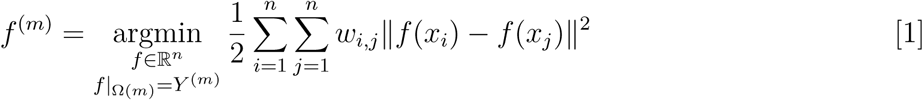

where *w*_*i,j*_ is a measure of the affinity between *x*_*i*_ and *x*_*j*_ that varies between 0 and 1, we compute it in 6.

This problem can be rewritten in terms of a matrix for each row (modality) *m*. First we introduce the Laplacian matrix *L*, given by

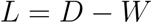

where *D* = diag(*d*_1_*, …, d*_*n*_) is the diagonal matrix with entries *d*_1_*, …, d*_*n*_ and 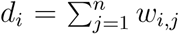 and *W* = (*w*_*i,j*_)_1*≤i,j≤n*_. With this, our optimization problem can be rewritten as

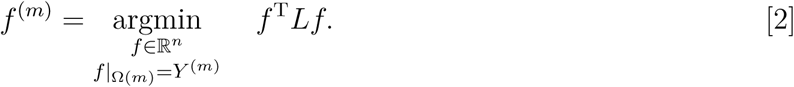

This is a quadratic optimization problem with linear constraints and can be solved by differentiating and setting the derivative to zero. The solution *f* ^(*m*)^, on the subsets Ω(*m*) and 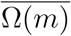, is given by

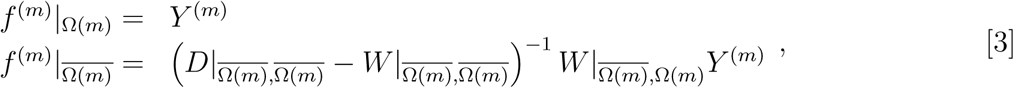

where we have adopted the notation of *M |*_*A,B*_ to denote the submatrix of *M* obtained by extracting the rows in the index set *A* and the columns in index set *B*.

The solution Eq. (3) can be rewritten as

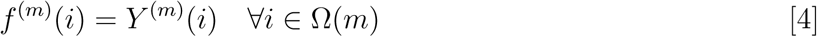

and

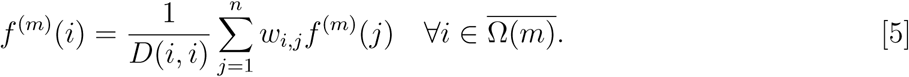

Note that Eq. (5) is an implicit equation satisfied by the values. To calculate the desired values of *f* ^(*m*)^, Eq. (3) must be used.

The above calculations are performed separately for each modality *m* = 1*, …, M*, each solution completing a different row in the overall measurement matrix.

### Computing the Affinity Matrix

Given a set of point (*x*_1_*, …, x*_*n*_), we compute the affinity matrix *W* = (*w*_*i,j*_) with

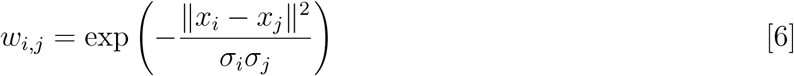

where the scaling parameters *σ*_*i*_ and *σ*_*j*_ are defined as 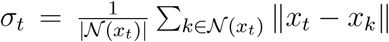 for every *t* = 1*, …, n*, where 𝒩 (*x*_*t*_) represents the neighborhood composed of the *|* 𝒩 (*x*_*t*_) *|* = 10 closest points of *x*_*t*_ (2). The norms here are Euclidean norms (3). This gives us a proximity scale that varies across the manifold if the sampling of the points is not uniform.

### K-Fold Cross Validation

To evaluate the accuracy of the reconstruction obtained by data fusion we employ K-fold cross validation (4). This approach estimates the expected prediction error. The idea is to predict the value of labels on a subset of the labeled data points and compare the prediction to the known labels.

Let us consider labeled data points *x|*_Ω(*m*)_ with labels *Y* ^(*m*)^ and unlabeled data points *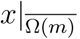.*The labeled data points can be uniformly assigned to *K* bins randomly using an indexing function κ: Ω(m) → {1, …, K}. For each value *k* of *κ*, we thus have the index set *κ*^*-*^1(k) = {i ∈ Ω(m): \κ(i) = k}, a subset of Ω(*m*). We can thus define reduced set of labeled measurements Ω_*k*_(*m*) = Ω(*m*) \*κ*^*-*1^(*k*), with the corresponding set of unlabeled measurements 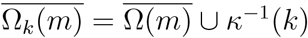.

For *i ∈* Ω(*m*), we denote by *f* ^(*m*)*,-κ*(*i*)^(*i*) the label prediction on data point *i* where Ω_*κ*(*i*)_(*m*) and 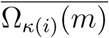 are used as the set of labeled and unlabeled points, respectively. Since 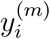 is known, we can evaluate the performance of our method by comparing the prediction *f* ^(*m*),-*κ*(*i*)^(*i*) to the known value 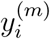. Repeating this for all *i ∈* Ω(*m*), we calculate the average absolute error normalized by the range for each modality *m*

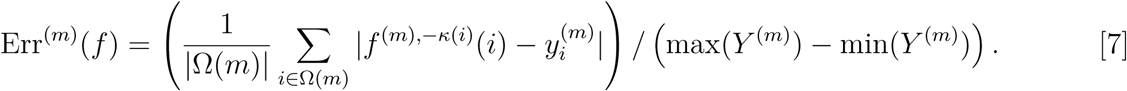

## 2. Illustrative example

To demonstrate the efficiency of this technique as an interpolation method on a non-linear manifold, we first consider a toy example with a non-linear 1-dimensional trajectory embedded in 3-dimensional space. Specifically, we have

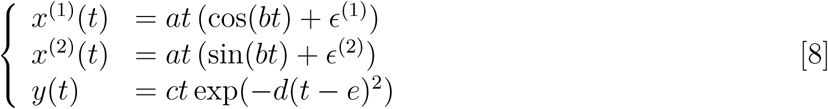

where, *a*, *b*, *c*, *d*, *e* are constants, *∈*^(1)^ and *∈*^(2)^ are Gaussian noise sources and *t* is a real-valued parameter. The set of points (*x*^(1)^(*t*)*, x*^(2)^(*t*)) forms a 1-dimensional non-linear manifold embedded in the 2-dimensional space as it is parameterized by *t*. These points correspond to the embryo morphology in our toy example. In the absence of noise, this mapping from *t* to the 2D plane can be inverted as 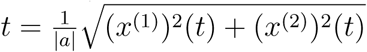. The signal *y*(*t*) is a smooth function of *t* and is thus a smooth function of (*x*^(1)^ (*t*)*, x*^(2)^ (*t*)) by composition. In this example, *y* corresponds to the target modality that we would like to estimate.

To reproduce the setting of data fusion with three modalities, ((*x*^(1)^*, x*^(2)^)*, t, y*), we consider the following situation. Suppose that one can acquire a set of labeled points, i.e. a set of *l* triplets, ((*x*^(1)^(*t*_1_)*, x*^(2)^(*t*_1_))*, y*(*t*_1_))*, …,* ((*x*^(1)^(*t*_*l*_)*, x*^(2)^(*t*_*l*_))*, y*(*t*_*l*_)) and a set of *u* unlabeled, but timestamped, points, ((*x*^(1)^(*t*_*l*+1_)*, x*^(2)^(*t*_*l*+1_))*, t*_*l*+1_)*, …,* ((*x*^(1)^(*t*_*l*+*u*_)*, x*^(2)^(*t*_*l*+*u*_))*, t*_*l*+*u*_), as shown in Fig. 1A and B. We therefore have Ω = *{*1*, …, l}* and 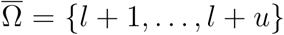 and *n* = *l* + *u* (since we only consider *y* as labels, and hence have only one modality to fill, we have suppressed the superscript (*m*)). The pairwise similarity measures *w*_*i,j*_ are computed using Euclidean norm between pairs of data points (*x*^(1)^(*t*_*i*_)*, x*^(2)^(*t*_*i*_)) and (*x*^(1)^(*t*_*j*_)*, x*^(2)^(*t*_*j*_)) as presented in 6. Then using the framework presented above, in particular equations Eq. (4) and Eq. (5), it is possible to estimate *y* = *f* (*x*) on the set of unlabeled data points using the harmonic extension algorithm. The results are shown in Fig. 1C. We then directly obtain *y* as a function of *t* by composition using the known time stamps (*t*_*l*+1_*, …, t*_*l*+*u*_).

For this example, we now quantify the evolution of the error as a function of the number of unlabeled samples. The constants used were *a* = 1, *b* = 0.1, *c* = 1, *d* = 0.005, *e* = 65, *l* = 120, and *u* = 300 while *t* varies from 1 to 100 and *∈*^(1)^ and *∈*^(2)^ are sampled according to centered Gaussian distributions with standard deviation 2.02. Using a K-fold validation strategy on the labeled samples, we characterize the accuracy of the reconstruction using the normalized absolute error as described by equation Eq. (7) (Fig. S1A). The results are shown on Fig. S1B. We found that with no additional data points, the absolute error was on average 1.05% of the signal range, and decreased uniformly to 0.53% when considering the problem with an additional 300 unlabeled data points.

## 3. Experimental Data Sets

### Data Sets description

To explore the capabilities of our data fusion strategy, we considered 11 different datasets, each of which consisting of a set of images, obtained either as live movies or fixed samples, and containing a channel recording the spatial distribution of nuclei.

### Live Movies

Nikon A1-RS confocal microscope with a 60x Plan-Apo oil objective was used to obtain time-lapse movies of Drosophila embryonic development. Live imaging of Histone-RFP embryos was used to visualize nuclei. A total of 7 movies were obtained, with a time resolution of 30 seconds per frame. All the movies start during nuclear cycle 14 (about 2.5 hr after fertilization) and end after about 20 min after gastrulation starts (about 3.3 hr after fertilization). The number of frames for each movie is 156, 67, 53, 55, 50, 86, and 90.

### Fixed samples

In our experiments, 4 datasets were acquired to visualize nuclei, protein expression of dpERK, Twist, and Dorsal, and mRNA expression of ind and rho. Immunostaining and fluorescent in situ hybridization protocols were used as described before (5). DAPI (1:10,000; Vector laboratories) was used to visualize nuclei. Rabbit anti-dpERK (1:100; Cell Signaling), mouse anti-Dorsal (1:100; DSHB), rat anti-Twist (1:1000; gift from Eric Wieschaus, Princeton University), sheep anti-digoxigenin (1:125; Roche), and mouse anti-biotin (1:125; Jackson Immunoresearch) were used as primary antibodies. Alexa Fluor conjugates (1:500; Invitrogen) were used as secondary antibodies. Stained embryos were imaged using Nikon A1-RS confocal microscope with a 60x Plan-Apo oil objective. Embryos were mounted in a microfluidic device for end-on imaging, as described previously (5, 6). All the images were taken at *∼*90 *μ*m from the posterior pole of the embryo.

### Dataset 1

Stained with rabbit anti-dpERK and rat anti-Twist antibodies. Number of samples: 108.

### Dataset 2

Stained with mouse anti-Dorsal antibody, rabbit anti-dpERK antibody, and ind-DIG probe. Number of samples: 59.

### Dataset 3

Stained with ind-biotin probe, rho-DIG probe, and rabbit-dpERK antibody. Number of samples: 58.

### Dataset 4

Stained with rat anti-Twist antibody, ind-biotin probe, and rho-DIG probe. Number of samples: 30.

### Preprocessing Steps

In order to compare the images of morphology (nuclei channel) among all the datasets, we used a certain number of preprocessing steps to overcome measurement-dependent variability.

### Image Rotation

Images were oriented so that the highest Dorsal signal is located at the ventral-most point. Some late-stages embryos were oriented manually.

### Image Resizing

The images were resized and cropped such that the embryo would occupy 80% of the image. This was achieved by thresholding the images and computing their bounding box, then resizing and translating the image so that the bounding box occupied the square stretching from 10% to 90% of the image width, both horizontally and vertically. The images were resized to 100 by 100 pixels.

### Intensity Renormalization

Since there is some local variation of image intensity that is due to the device used to acquire data and the configuration of the microscope, we wish to remove this from the images before comparing them. This is done by computing a local average using a Gaussian kernel, and then renormalizing the image by that value. Specifically, given an image *x*, we calculate its local average *y*(*u*) = *x * ϕ*(*u*), where 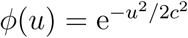 for some kernel width *c* > 0 and *\** denotes convolution. Then the renormalized image is given, for each pixel *u*, by

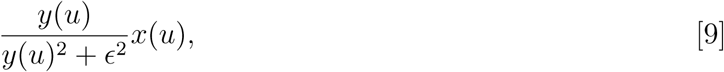

where *∈* > 0 is some threshold that prevents the renormalization from exploding when the image is locally close to zero.

### Contrast Increase

Another consequence of the experimental setup and the configuration of the microscope is that the contrast differs between measurements. As a result, the range of pixel intensities is not the same for different images of embryos at the same developmental stage. To fix this, we pass the pixel intensities through a logistic function with two parameters. Given an input image *x*, it is adjusted to give, for each pixel *u*,

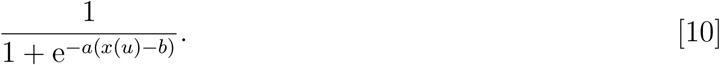

### Scattering Transform

To compare images without being sensitive to small translations or deformations, we applied the scattering transform (7) and compared the resulting transform vectors. Differences between images in the scattering space are meant to reflect changes in the underlying morphology as opposed to variation due to the imaging method.

The scattering transform of an image is a signal representation obtained by alternating wavelet decompositions and pointwise modulus operators. The resulting coefficients are locally invariant to translations and stable to deformations. The similarity of the signals can then be measured by computing the Euclidean distance of their scattering transforms. These transforms have enjoyed significant success in classification of images (8, 9), but also time series (10), since these tasks often requires insensitivity to translation and deformation.

For our embryo images, we have found that second-order scattering coefficients with an averaging scale of 64 pixels allow for the proper amount of invariance. These are computed using the ScatNet toolbox (11).

The result is a vector of dimension 784 for each image. The point clouds corresponding to each of the 11 datasets were centered separately.

### Computing the affinity matrix

The affinity matrix was computed according to the definition of the Gaussian kernel with an adaptive scaling factor, as described in 6. The result is shown on Fig. S3.

### Outlier filtering

Even after applying all the previous steps of image preprocessing some data points were considerably away from the main manifold. First we calculated the closest neighbor of each point. We then kept the points whose closest neighbor distance was less than 3 times the median minimal neighbor distance.

### Results

To fuse the channels of the various datasets described in 3, we applied the harmonic extension algorithm described in 1. The data points are preprocessed images of nuclei spatial distribution using the steps presented in 3 to 3 with parameter values as shown in Table S1 and the modalities correspond to the images obtained from the various fluorescent reporters described in section 3 of the SI appendix. The corresponding low-dimensional manifold on which the data points *x*_*i*_ lie is shown on Fig. S4.

We denote by *x*_*i*_ the preprocessed image *i* of the nuclei channel, it is an element of ℝ ^784^ as the result of scattering transformation applied to a 100 *×* 100 image. We denote by 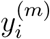 the image corresponding to modality associated to and by 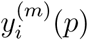 its restriction to the pixel *p*, 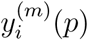 is an element of ℝ.

For each of the 10000 pixels in the live movie, there are 5 modalities that we would like to fuse: dpERK (*m* = 1), Twist (*m* = 2), Dorsal (*m* = 3), ind (*m* = 4), rho (*m* = 5). We therefore solve the data fusion problem for each pixel and each modality, leading to 50000 semi-supervised learning solutions. The combination of labeled and unlabeled datasets is described on Table S2.

To evaluate the accuracy of the method, we computed the cross-validation error as described by equation Eq. (7) for each pixel and averaged over the entire images. The range of the signal is calculated based on the entire images. Table S3 shows the results for each of the datasets where the number of unlabeled data points is 309 (the number of live movie frames), the unlabeled data points are chosen randomly among the live movie frames and the fixed images from the 4 datasets which serve as unlabeled data points. For each of the datasets, we chose *K* such the number of points within each bin was about 20. We have *K* = 6, 3, 3, 2 for datasets 1 to 4 respectively.

### Coloring movies

The resulting fused dataset led to the construction of a multimodal movie, where a different color was attributed to each modality and the RGB values were added in the resulting movie. We used the set of colors presented on Table S4.

---

**Supplementary Files 1:** This compressed folder contains all the raw images organized into folders: movie1-7, and data_set1-4. The folders movie1-7 correspond to the live movies, while the folders data_set1-4 correspond to stained fixed images. The description of the datasets can be found within each folder or in section 3 of the SI appendix.

**Supplementary Files 2:** Colored movie. The compressed folder contains frames of a colored movie, with, in the main folder, all the channels combined, nuclei (gray), dpERK (red), Dl (pink), rho (yellow), ind (blue), Twi (green), using the color map presented in Table S4. The number of each image corresponds to a time step, and they have been extracted from movie 5. The subfolder “grayscale” contains the actual value of the variables in separate frames, the name of each image indicates the variable considered, while the number of each image corresponds to the time step.

**Supplementary Software:** The compressed folder contains the MATLAB library implementing the data fusion algorithm on an illustrative example and on experimental cross-sections of developing fly embryos. The file README.md contains instructions on how to run the code. The main scripts are found in the examples folder and consist of:

- spiral_example.m: implements the data fusion algorithm on a non-linear 1-dimensional trajectory in a 3-dimensional space, nicknamed “spiral”.
- spiral_example_cross_validation.m: implements the K-fold cross-validation on the “spiral”.
- experimental_dataset.m: implements the data fusion on the experimental datasets leading to a multimodal movie containing the spatio-temporal dynamics of 5 chemical species on top of morphological changes.
- experimental_dataset_cross_validation.m: implements the K-fold cross validation on the experimental datasets.

**Table S1.** Values of the parameters for intensity renormalization (Section 3) and contrast increase (Section 3) for each of the experimental datasets.

|  | Live Movies | Dataset 1 | Dataset 2 | Dataset 3 | Dataset 4 |
| --- | --- | --- | --- | --- | --- |
| $\epsilon$ | 0.01 | 0.01 | 0.01 | 0.05 | 0.01 |
| $c$ | 10 | 10 | 10 | 5 | 10 |
| $a$ | 10 | 10 | 10 | 15 | 10 |
| $b$ | 0.8 | 1.0 | 1.0 | 0.5 | 1.0 |

**Table S2.** Distribution of the datasets into labeled and unlabeled sets depending on the modality. We refer to *Ω(m*) as the set of labeled datapoints, while 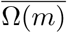 is the set of unlabeled data points for the *m*th modality.

|  | Live Movies | Dataset 1 | Dataset 2 | Dataset 3 | Dataset 4 |
| --- | --- | --- | --- | --- | --- |
| $m = 1$<br>dpERK | $\overline{\Omega(1)}$ | $\Omega(1)$ | $\Omega(1)$ | $\Omega(1)$ | $\overline{\Omega(1)}$ |
| $m = 2$<br>Twist | $\overline{\Omega(2)}$ | $\Omega(2)$ | $\overline{\Omega(2)}$ | $\overline{\Omega(2)}$ | $\Omega(2)$ |
| $m = 3$<br>Dorsal | $\overline{\Omega(3)}$ | $\overline{\Omega(3)}$ | $\Omega(3)$ | $\overline{\Omega(3)}$ | $\overline{\Omega(3)}$ |
| $m = 4$<br>ind | $\overline{\Omega(4)}$ | $\overline{\Omega(4)}$ | $\Omega(4)$ | $\Omega(4)$ | $\Omega(4)$ |
| $m = 5$<br>rho | $\overline{\Omega(5)}$ | $\overline{\Omega(5)}$ | $\overline{\Omega(5)}$ | $\Omega(5)$ | $\Omega(5)$ |

**Table S3.**
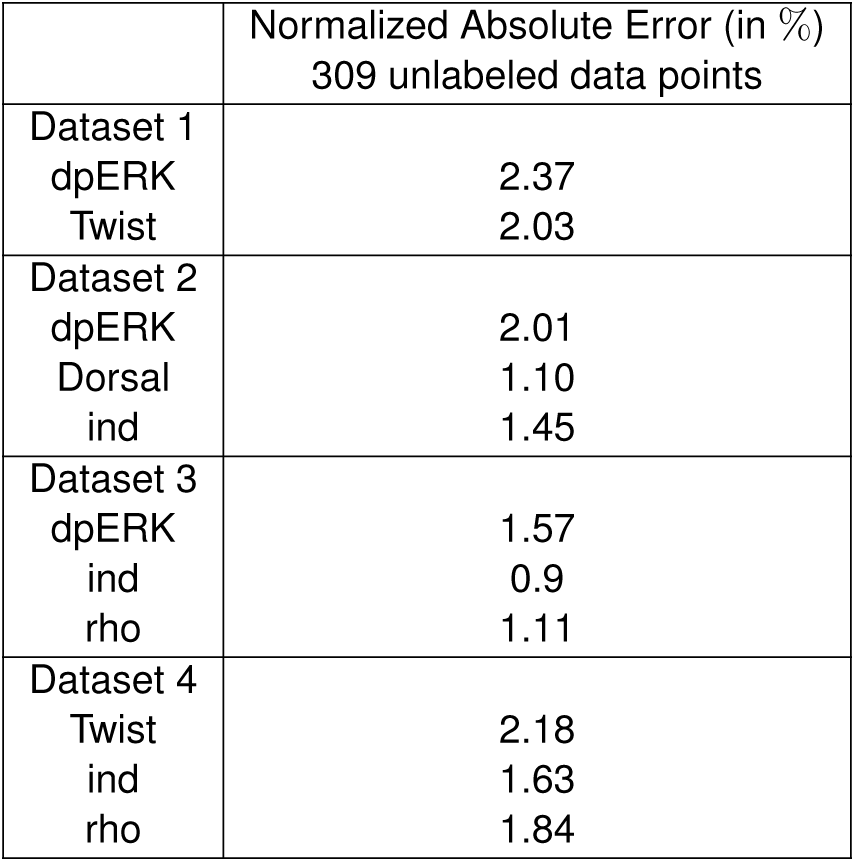
Normalized Absolute Error obtained by K-fold cross-validation for each modality of each dataset. In each case, we performed 10 repetitions, where the labeled samples are distributed randomly among the K bins, and the 309 unlabeled data points are chosen randomly. The error is then averaged over 10 repetitions. More details about the Normalized Absolute Error can be found in section 1 of the SI appendix and about how it is computed on the real datasets in section 3 of the SI appendix.

|  | Normalized Absolute Error (in %)<br>309 unlabeled data points |
| --- | --- |
| Dataset 1 |  |
| dpERK | 2.37 |
| Twist | 2.03 |
| Dataset 2 |  |
| dpERK | 2.01 |
| Dorsal | 1.10 |
| ind | 1.45 |
| Dataset 3 |  |
| dpERK | 1.57 |
| ind | 0.9 |
| rho | 1.11 |
| Dataset 4 |  |
| Twist | 2.18 |
| ind | 1.63 |
| rho | 1.84 |

**Table S4.** Color scheme used to color the final movie.

|  | R | G | B |
| --- | --- | --- | --- |
| dpERK | 213 | 36 | 2 |
| Twist | 92 | 188 | 146 |
| Dorsal | 229 | 195 | 207 |
| ind | 103 | 139 | 191 |
| rho | 225 | 223 | 75 |

**Fig. S1.**
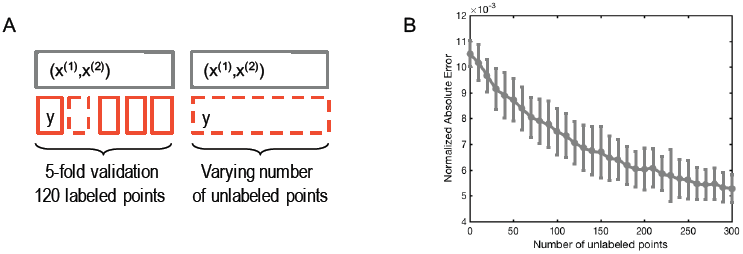
K-fold cross validation on the illustrative example A) Setting with *K* = 5, there are 120 labeled points and the number of unlabeled points varies from 0 to 300. B) The normalized absolute error as a function of the number of unlabeled points. There are 100 repetitions for each number of unlabeled data points.

**Fig. S2.**
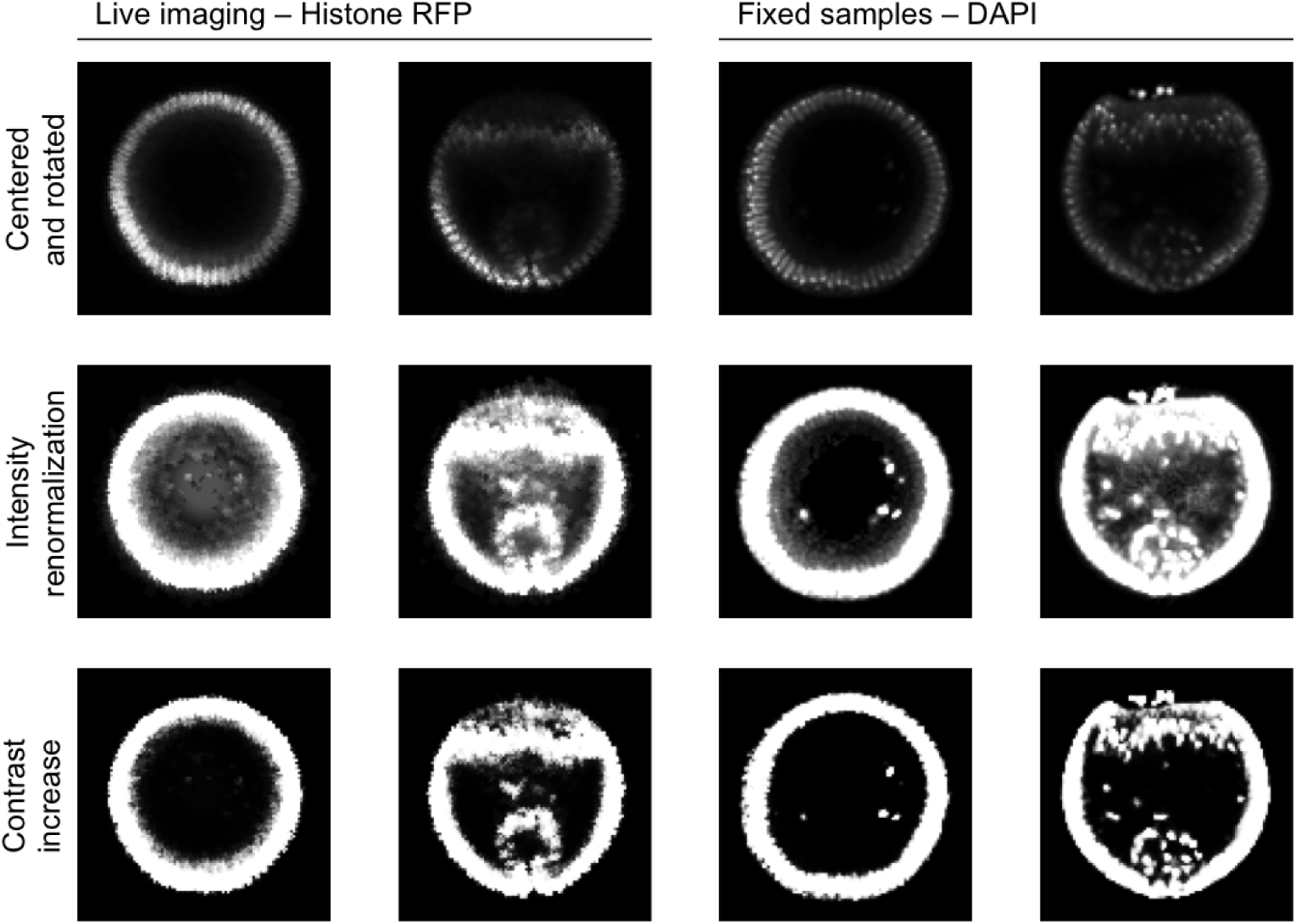
Illustration of the image preprocessing steps applied on the nuclei channel and described in section S2 of the SI appendix. The first line shows images resulting from rotation and centering steps. The second line shows images resulting from intensity renormalization. The third line shows images resulting from contrast increase. The first two columns show early and later stages from movie frames stained with Histone-RFP. The last two columns represent early and later stages from fixed samples stained with DAPI.

**Fig. S3.**
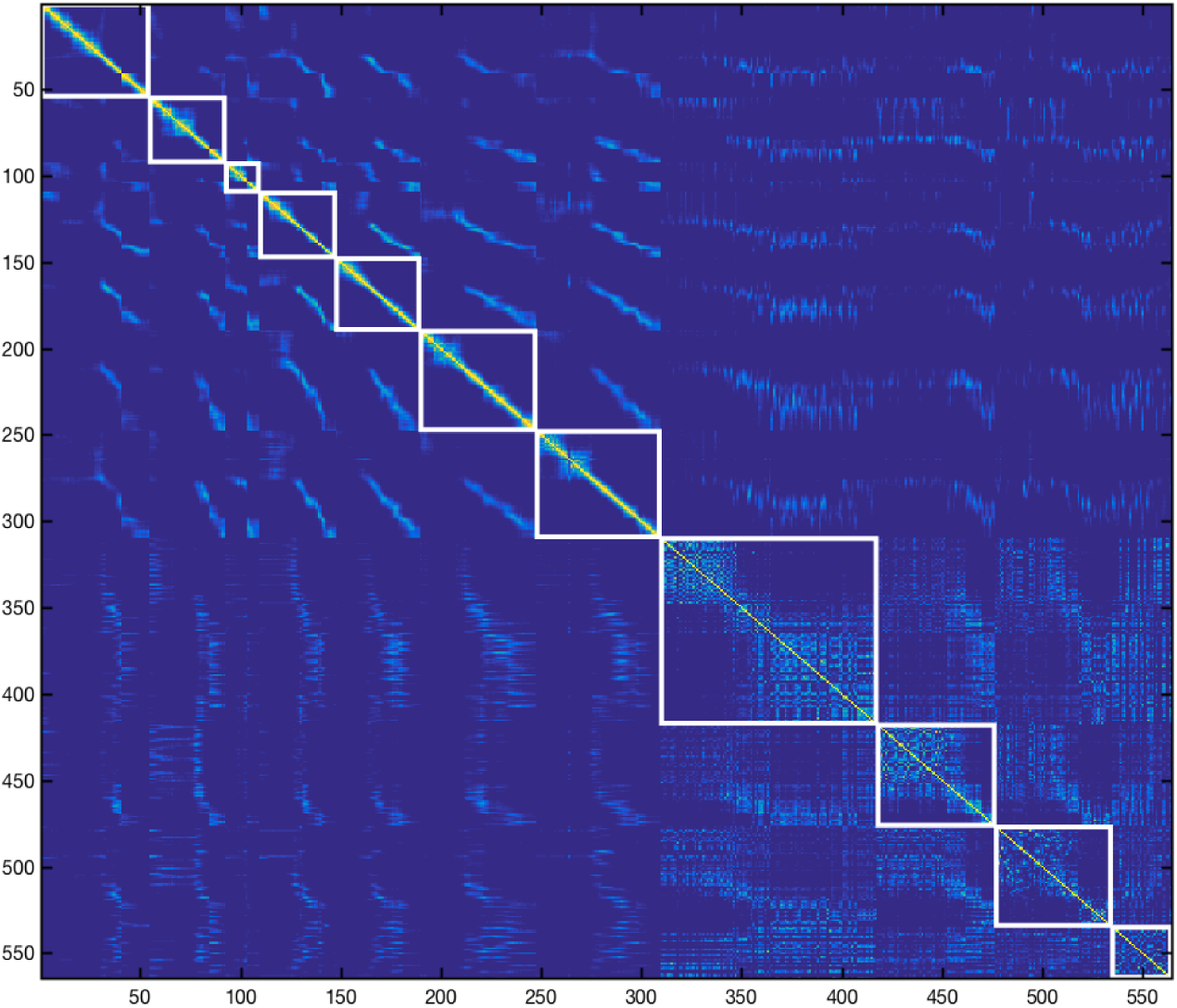
The Affinity Matrix *W* = (*w*_*i,j*_) obtained by comparing images as described by equation 6 in SI appendix is shown as a heatmap. The white squares identify each of the 11 datasets. The first 7 correspond to live movies, the last 4 correspond to the datasets of fixed images.

**Fig. S4.**
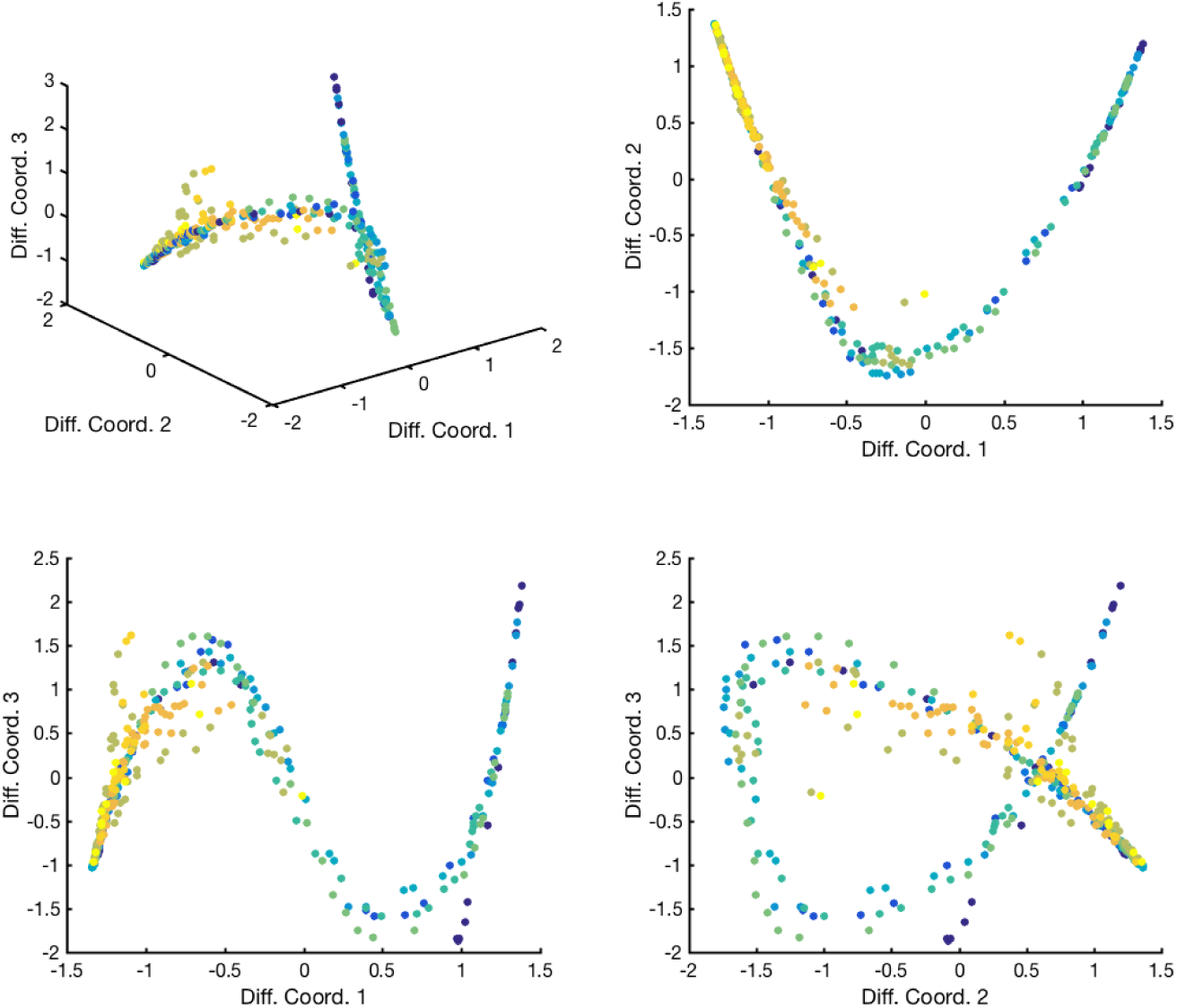
Low-dimensional embedding of the 11 datasets obtained by diffusion maps. Each dot is a point and each color is a different dataset. The top left panel shows the points obtained by embedding the points in the first three diffusion map coordinates. The top right panel shows the data points in the plane formed by the first two diffusion map coordinates, while the two bottom panels show the embedding in the planes obtained with the first and third (left) or second and third (right) diffusion map coordinates. Some outliers were filtered out for visualization purposes if their closest neighbor distance was at least twice the median closest neighbor distance, leading to a very well-defined 1-dimensional manifold.

